# Insight into mosquito GnRH-related neuropeptide receptor specificity revealed through analysis of naturally occurring and synthetic analogs of this neuropeptide family

**DOI:** 10.1101/699645

**Authors:** Azizia Wahedi, Gerd Gäde, Jean-Paul Paluzzi

**Author notes:** corresponding authors Prof. Jean-Paul Paluzzi, Department of Biology, York University, 4700 Keele Street, Toronto, Ontario, M3J1P3, Canada,; Prof. Gerd Gäde, Department of Biological Sciences, University of Cape Town, Private Bag, Rondebosch 7701, South Africa.

## Abstract

Adipokinetic hormone (AKH), corzazonin (CRZ) and the AKH/CRZ-related peptide (ACP) are peptides considered homologous to the vertebrate gonadotropin-releasing hormone (GnRH). All three *Aedes aegypti* GnRH-related neuropeptide receptors have been characterized and functionally deorphanized, which individually exhibit high specificity for their native ligands, which prompted us to investigate the contribution of ligand structures in conferring receptor specificity. In the current study, we designed a series of analogs based on the native ACP sequence in *A. aegypti* and screened them against the ACP receptor using a heterologous system to identify critical residues required for receptor activation. Specifically, analogs lacking the carboxy-terminal amidation, replacing aromatic residues, as well as truncated analogs were either completely inactive or had very low activities even at high concentration. The next most critical residues were the polar threonine in position 3 and the blocked amino-terminal pyroglutamate, with activity of the latter partially recovered using an alternatively blocked analog. ACP analogs with alanine substitutions at position 2 (valine), 5 (serine), 6 (arginine) and 7 (aspartic acid) positions were less detrimental as were replacements of charged residues. Interestingly, replacing asparagine with an alanine at position 9, creating a C-terminal WAA-amide, resulted in a 5-fold more active analog which may be useful as a lead superagonist compound. Similarly, we utilized this high-throughput approach against an *A. aegypti* AKH receptor (AKHR-IA) testing a number of mostly naturally-occurring AKH analogs from other insects to determine how substitutions of specific amino acids in the AKH ligand influences receptor activation. AKH analogs having single substitutions compared to the endogenous *A. aegypti* AKH revealed position 7 (serine) was well tolerated whereas changes to position 6 (proline) had pronounced effects, with receptor activity compromised nearly ten-fold. Substitution of position 3 (threonine) or analogs with combinations of substitutions were quite detrimental with a significant decrease in AKHR-IA activation. Interestingly, analogs with an asparagine residue at position seven displayed improved receptor activation compared to the native mosquito AKH. Collectively, these results advance our understanding of how two GnRH-related systems in *A. aegypti* sharing the most recent evolutionary origin sustain independence of function and signalling despite their relatively high degree of ligand and receptor homology.

## Introduction

In invertebrates there exist three neuropeptide systems of which the mature peptides as well as their cognate G protein-coupled receptors (GPCRs) are structurally similar, and collectively, they are related to the vertebrate gonadotropin releasing hormone (GnRH) system and suggested to form a large peptide superfamily (Gäde et al., 2011; Hansen et al., 2010; Li et al., 2016; Roch et al., 2011). These neuropeptide systems are called the adipokinetic hormone (AKH)/red pigment-concentrating hormone (RPCH) family, the corazonin (CRZ) family and the structurally intermediate adipokinetic hormone/corazonin-related peptide (ACP) family. AKHs are primarily or exclusively produced in neurosecretory cells of the corpus cardiacum, and a major function is mobilization of energy reserves stored in the fat body, thus providing an increase of the concentration of diacylglycerols, trehalose or proline for locomotory active phases in the haemolymph. In accordance with this function, AKH has been shown to activate the enzymes glycogen phosphorylase and triacylglycerol lipase and transcripts of the AKHR are most prominently found in fat body tissue (see reviews by (Gäde, 2004; Gäde and Auerswald, 2003; Marco and Gäde, 2019a). As with most endocrine regulatory peptides, additional functions such as inhibition of anabolic processes (protein and lipid syntheses), involvement in oxidative stress reactions and egg production inter alia are known making AKH a truly pleiotropic hormone (see reviews by (Kodrík, 2008; Kodrík et al., 2015). This is also true for CRZ which is mainly synthesized in neuroendocrine cells of the pars lateralis of the protocerebrum and released via the corpora cardiaca (Predel et al., 2007). Although originally described as a potent cardiostimulatory peptide (Veenstra, 1989), it does not generally fulfil this role but is known for many additional functions such as involvement in the (1) release of pre-ecdysis and ecdysis triggering hormones, (2) reduction of silk spinning rates in the silk moth, (3) pigmentation events (darkening) of the epidermis in locusts during gregarization and (4) regulation of caste identity in an ant species (Gospocic et al., 2017; Kim et al., 2004; Tanaka et al., 2002; Tawfik et al., 1999). The functional role of ACP is less clear. All previous studies had not found a clear-cut function for this peptide until work by Zhou and colleagues claimed that ACP in the cricket *Gryllus bimaculatus* regulates the concentration of carbohydrates and lipids in the haemolymph (Zhou et al., 2018).

Experiments in the current study have been conducted with the mosquito species *Aedes aegypti* for a number of reasons. Primarily because of being such an infamous disease vector for pathogens such as Yellow and Dengue fever, Chikungunya and, as latest addition, Zika arboviruses, all of which summarily are responsible for affecting approximately 50-100 million people per year, with the majority of these attributed to Dengue virus infection (Baud et al., 2017; Price et al., 2015; Saiz et al., 2016). Secondly, knowledge on the interaction of the ligands with their respective receptor are thought to be very helpful for drug research using specific GPCRs as targets for development of selective biorational insecticides (Audsley and Down, 2015; Hill et al., 2018; Verlinden et al., 2014). Another reason was that each of the three neuropeptide systems has already been partially investigated, thus, this study could build on previous research data. With respect to the peptides, these were first predicted from the genomic work on *A. aegypti* (Nene et al., 2007). The presence of mature AKH and corazonin, but not of ACP, was shown by direct mass profiling (Predel et al., 2010). The AKH and ACP precursors have been cloned (Kaufmann et al., 2009) as have one CRZ receptor (CRZR) and various variants of the AKHR and ACPR (Kaufmann et al., 2009; Oryan et al., 2018; Wahedi and Paluzzi, 2018). Each receptor is very selective and responds only to its cognate peptide, thus there are indeed three completely separate and independently working neuroendocrine systems active in this mosquito species.

With respect to the ACP system in *A. aegypti*, expression studies revealed transcripts of the major ACPR form as well as the ACP precursor were enriched during development in adults, specifically during day one and four after eclosion (Kaufmann et al., 2009; Wahedi and Paluzzi, 2018). On the tissue level, ACP precursor transcripts are mainly present in the head/thorax region (Kaufmann et al., 2009) and, more specifically, within the brain and thoracic ganglia (Wahedi and Paluzzi, 2018) and the ACPR transcripts are mainly detected in the central nervous system with significant enrichment in the abdominal ganglia, particularly in males (Wahedi and Paluzzi, 2018). The AKH precursor transcripts are present in head/thorax region of pupae and adult females and in the abdomen of males (Kaufmann et al., 2009). Expression of AKHR was shown in all life stages (Kaufmann et al., 2009), but similar to the ACPR and ACP transcripts, AKHR transcript variants were found to be significantly enriched in early adult stages (Oryan et al., 2018). On the tissue level, highest expression in adults was found in thorax and abdomen with low levels detected in ovaries (Kaufmann et al., 2009). Similar expression profiles were determined recently by Oryan et al. (2018) using RT-qPCR: expression in head, thoracic ganglia, accessory reproductive tissues and carcass of adult females as well as in abdominal ganglia and specific enrichment in the carcass and reproductive organs of adult males. Similarly, CRZR was mostly expressed during later development, including pupal and adult stages. In terms of tissue expression profiles, this was examined in adult stages where CRZR was found in the heads and thoracic ganglia of both sexes as well as in ovary, abdominal ganglia and Malpighian tubules in females and in the testes and carcass of males (Oryan et al., 2018).

Thus, although quite a substantial body of information on the three signalling systems in *A. aegypti* is known, we lack knowledge of how the various ligands interact with their cognate receptor. For instance, which amino acid residue in the ligand is specifically important for this interaction? By replacing each residue successively with simple amino acids such as glycine or alanine one can probe into the importance of amino acid side chains and their characteristics in relation to facilitating receptor specificity. Such structure-activity studies have been conducted on the AKH system in a number of insects and a crustacean either by measuring physiological actions *in vivo* (Gäde and Hayes, 1995; Ziegler et al., 1998) or in a cellular mammalian expression system *in vitro* (Caers et al., 2012; Marchal et al., 2018; Marco et al., 2017). In conjunction with nuclear magnetic resonance (NMR) data on the secondary structure of AKHs (Nair et al., 2001; Zubrzycki and Gäde, 1994) and the knowledge of the receptor sequence, molecular dynamic methods can devise models how the ligand is interacting with its receptor (Jackson et al., 2018; Mugumbate et al., 2013).

Since no such data is available for any ACP system, the objective of the current study was to fill this knowledge gap by determining critical amino acids of the ACP neuropeptide necessary for activation of its cognate receptor, ACPR. We therefore designed a series of analogs based on the endogenous *A. aegypti* ACP sequence and screened them using an *in vitro* receptor assay system to determine the crucial residues for ACPR activation. Given the closer evolutionary relationship between the ACP and AKH systems in arthropods (Hansen et al., 2010; Hauser and Grimmelikhuijzen, 2014; Tian et al., 2016; Zandawala et al., 2018), a second aim of the current study was to advance our knowledge and understanding of critical residues and properties of uniquely positioned amino acids necessary for the specificity of the AKH ligand to its receptor, AKHR-IA. Here, we took advantage of naturally occurring AKHs from other insects to examine amino acid substitutions and determine the consequences onto AKHR-IA activation using the same *in vitro* heterologous assay. Collectively, determining indispensable amino acid residues of these two GnRH-related mosquito neuropeptides that are necessary for activation of their prospective receptors will help clarify how these evolutionarily-related systems uphold specific signalling networks avoiding cross activation onto the structurally related, but functionally distinct signalling system.

## Materials and Methods

### ACP and AKH Receptor Expression Construct Preparation

A mammalian expression construct containing the ACPR-I ORF (hereafter denoted as ACPR) with Kozak translation initiation sequence had been previously prepared in pcDNA3.1^+^ (Wahedi and Paluzzi, 2018); however, this earlier study revealed that coupling of the ACPR with calcium mobilization in the heterologous expression system was not very strong, even following application of high concentrations of the native ACP ligand. Therefore, to improve the bioluminescent signal linked to calcium mobilization following receptor activation, which was required in order to provide greater signal to noise ratio when testing ACP analogs with substitutions leading to intermediate levels of activation of the ACPR, we created a new mammalian expression construct using the dual promoter vector, pBudCE4.1. Specifically, we utilized a cloning strategy as reported previously (Paluzzi et al., 2015) whereby the ACPR was inserted into the multiple cloning site (MCS) downstream of the CMV promoter in pBudCE4.1 using the sense primer *Sal*I-KozakACPR, 5’-G<u>GTCGAC</u>GCCACCATGTATCTTTCGG-3’ and anti-sense primer *Xba*I-ACPR, 5’-<u>TCTAGA</u>TTATCATCGCCAGCCACC-3’, which directionally inserted the ACPR within the MCS downstream of the CMV promoter. Secondly, the murine homolog (Gα15) of the human promiscuous G protein, Gα16, which indiscriminately couples a wide variety of GPCRs to calcium signaling (Offermanns and Simon, 1995) was inserted into the MCS downstream of the EF1-α promoter in pBudCE4.1 using the sense primer *Not*I-Gα15, 5’-GCGGCCGCCACCATGGCCCGGTCCCTGAC-3’ and anti-sense primer *Bgl*II-Gα15, 5’-AGATCTTCACAGCAGGTTGATCTCGTCCAG-3’, which directionally inserted the murine Gα15 within the MCS downstream of the EF1-α promoter.

The AKHR mammalian expression construct used in the current study, specifically AKHR-IA which was the receptor isoform that exhibited the greatest sensitivity to its native AKH ligand, was prepared previously in pcDNA3.1^+^ (Oryan et al., 2018) and coupled strongly with calcium signaling in the heterologous system so there was no need utilize the dual promoter vector containing the promiscuous Gα15 as was necessary for ACPR. For each construct, an overnight liquid culture of clonal recombinant bacteria containing the appropriate plasmid construct was grown in antibiotic-containing LB media and used to isolate midiprep DNA using the PureLink Midiprep kit (Life Technologies, Burlington, ON). Midiprep samples were then quantified and sent for sequencing as described previously (Oryan et al., 2018; Wahedi and Paluzzi, 2018).

### Cell culture, transfections, and calcium bioluminescence reporter assay

Functional activation of the AedaeACPR and AedaeAKHR-IA receptors was carried out following a previously described mammalian cell culture system involving a Chinese hamster ovary (CHO)-K1 cell line stably expressing aequorin (Paluzzi et al., 2012). Cells were grown in DMEM:F12 media containing 10% heat-inactivated fetal bovine serum (Wisent, St. Bruno, QC), 200μg/mL Geneticin, 1x antimycotic-antibiotic to approximately 90% confluency, and were transiently transfected using either Lipofectamine LTX with Plus Reagent or Lipofectamine 3000 transfection systems (Invitrogen, Burlington, ON) following manufacturer recommended DNA to transfection reagent ratios. Cells were detached from the culture flasks at 48-hours post-transfection using 5mM EDTA in Dulbecco’s PBS. Cells were then prepared for the receptor functional assay following a procedure described previously (Wahedi and Paluzzi, 2018). Stocks of peptide analogs of ACP and AKH were used for preparation of serial dilutions all of which were prepared in assay media (0.1% BSA in DMEM:F12) and loaded in quadruplicates into 96-well white luminescence plates (Greiner Bio-One, Germany). Cells were loaded with an automatic injector into each well of the plate containing different peptide analogs at various concentrations as well as negative control (assay media alone) and positive control wells (50µM ATP). Immediately after injection of the cells, luminescence was measured for 20 seconds using a Synergy 2 Multi-Mode Microplate Reader (BioTek, Winooski, VT, USA). Calculations, including determination of EC_50_ values, were conducted in GraphPad Prism 7.02 (GraphPad Software, San Diego, USA) from dose-response curves from 3-4 independent biological replicates.

### Synthetic Peptides and Analogs

Peptide analogs based on the native ACP sequence were designed and synthesized by Pepmic Co., Ltd. (Suzhou, China) at a purity of over 90%. All synthetic peptides were initially prepared as stock solutions at a concentration of 1mM in dimethyl sulfoxide. All stocks of arthropod AKH analogs were prepared similarly to the ACP synthetic peptide analogs and were commercially synthesized by either Pepmic Co., Ltd. (Suzhou, China) or by Synpeptide Co. (Shanghai, China) at a purity of over 90%. Specific sequence information for each ACP and AKH peptide analog used in this study is provided in Tables 1 and 2, respectively.

**Table 1.**
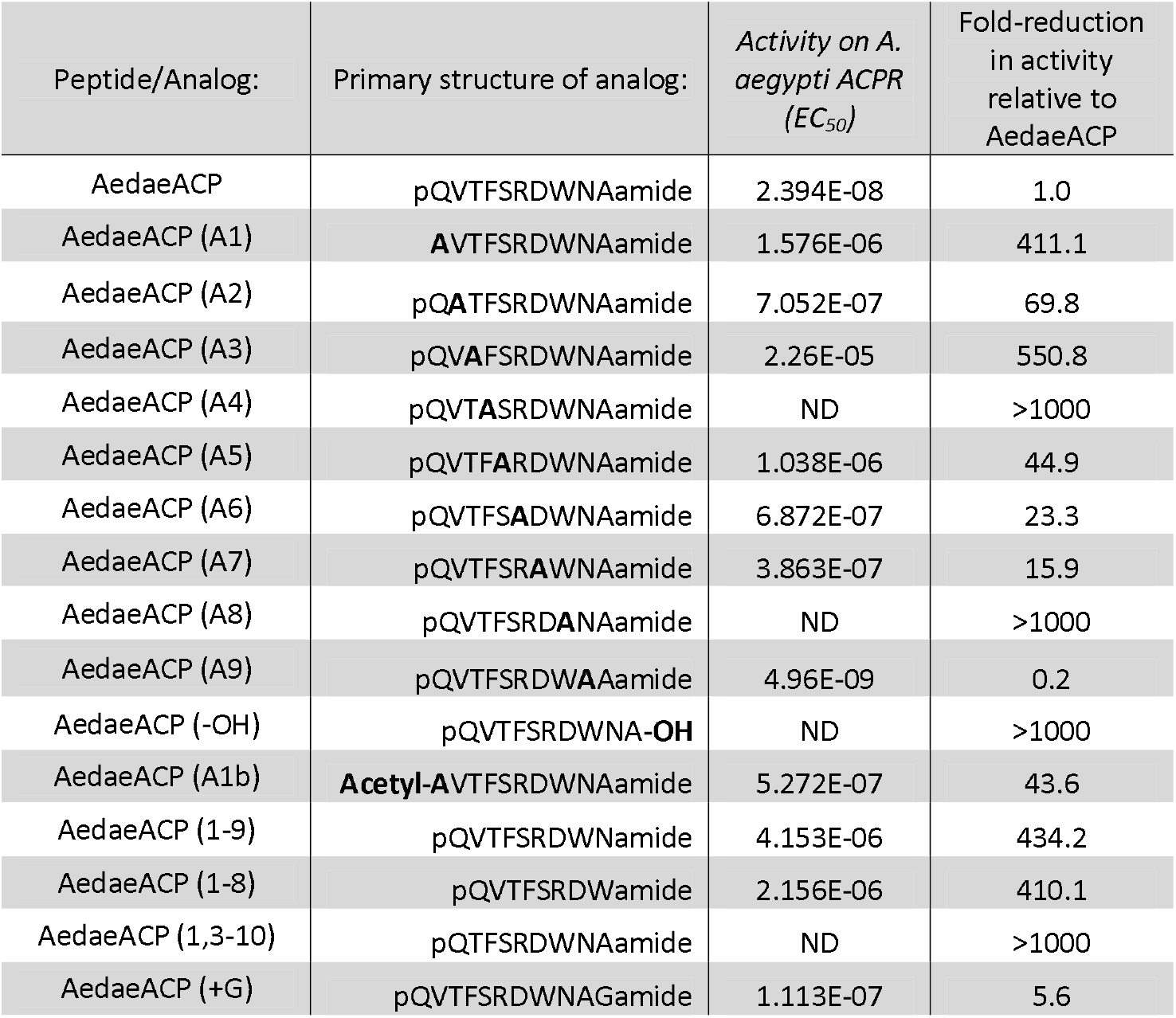
Primary structure of the synthetic ACP analogs designed and tested on the *A. aegypti* ACP receptor (AedaeACPR). Using heterologous expression of ACPR, the half maximal effective concentration for each of the analogs was determined as well as the corresponding reduction in activity, highlighting how a particular residue substitution or other modification is tolerated in this system. ND = represents no determined EC_50_ value due to minimal or no detectable activity of the analog when tested up to 10μM.

**Table 2.**
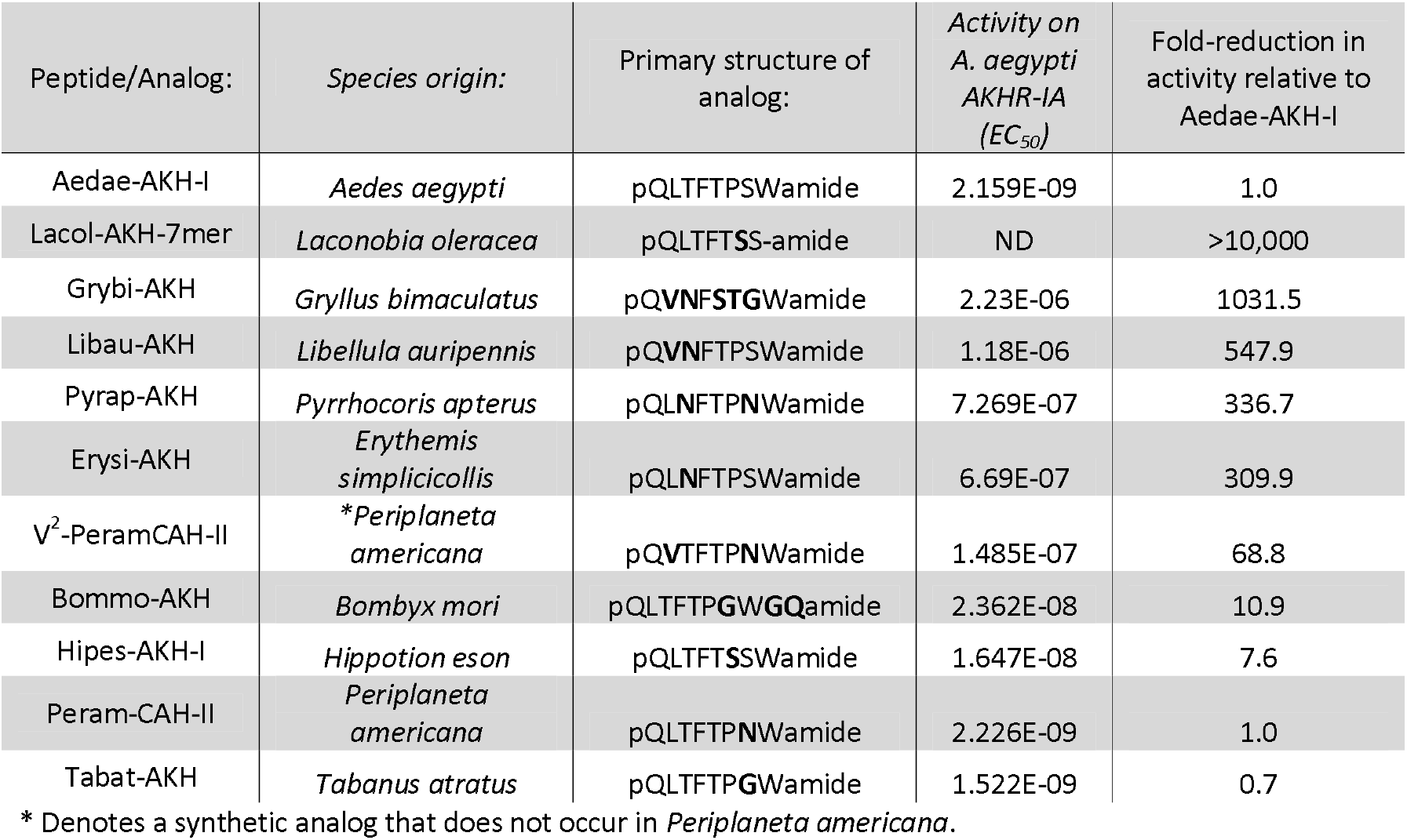
Primary structure of select naturally occurring AKH analogs from insects and their activity on the *A. aegypti* AKHR-IA receptor (AKHR-IA). Using heterologous expression of AKHR-IA, the half maximal effective concentration for each of the analogs was determined as well as the corresponding reduction in activity, highlighting how a particular single residue substitution, a combination of substitutions or other modifications are tolerated in this system. ND = represents no determined EC_50_ value due to minimal or no detectable activity of the analog when tested up to 10μM.

## Results

### ACP synthetic analog activity on mosquito ACPR

Analogs of native ACP with single alanine substitutions at each amino acid were synthesized and evaluated by examining their functional activation of the *A. aegypti* ACP receptor (ACPR) expressed using a heterologous system. Carboxy-terminal amidation of ACP was found to be critical for bioactivity since the free acid ACP analog elicited no activation of ACPR (Figure 1A) detectable up to the highest tested concentration of 10uM (Table 1). In addition, analogs replacing either of the two aromatic residues of ACP, including the phenylalanine in the fourth position or tryptophan in the eighth position, were similarly inactive on the ACPR (Figure 1A) with no detectable response by any of the tested concentrations (Table 1). Further examination of single residue alanine substitutions revealed that the next most critical residue for retaining ACPR functional activation was the polar threonine in the third position, which upon replacement with alanine resulted in a ~551-fold reduced activity compared to native ACP (Figure 1A, Table 1). A comparable reduction of activity, approximately 411-fold, was measured with an ACP analog that had the first residue, the blocked pyroglutamate in the native ACP, substituted by alanine. If the substitution, however, was also made with a blocked compound, N-Acetyl-alanine instead of pGlu, the analog was 10-times more active as the non-blocked one, thus the reduction in activity was only about 44-fold (Figure 1A). Interestingly, substitutions with analogs containing alanine at the second (valine), fifth (serine), sixth (arginine) and seventh (aspartic acid) positions, had noticeably less impact on the reduction of activity (Figure 1B), by ~70-fold, ~45-fold, ~23-fold, and ~16-fold, respectively. Unexpectedly, the analog containing an alanine substitution for the asparagine in position nine resulted in a more potent ACPR activation, leading to a ~5-fold greater activity compared to the native ACP peptide (Figure 1B). C-terminally truncated analogs, which lacked either the naturally occurring alanine residue in the tenth residue position or the last two C-terminal residues including asparagine and alanine in positions nine and ten, were almost completely inactive, demonstrating ~434-fold and ~410-fold reduced ACPR activation compared to native ACP (Figure 1C). A C-terminally extended analog ending with a glycine on the C-terminus elicited a small decrease of activity by about ~6-fold (Figure 1C). Lastly, the internally truncated ACP analog lacking a valine in the second position also was without any effect, similar to the two aromatic residue substituted analogs or the non-amidated ACP analog (Figure 1C).

**Figure 1.**
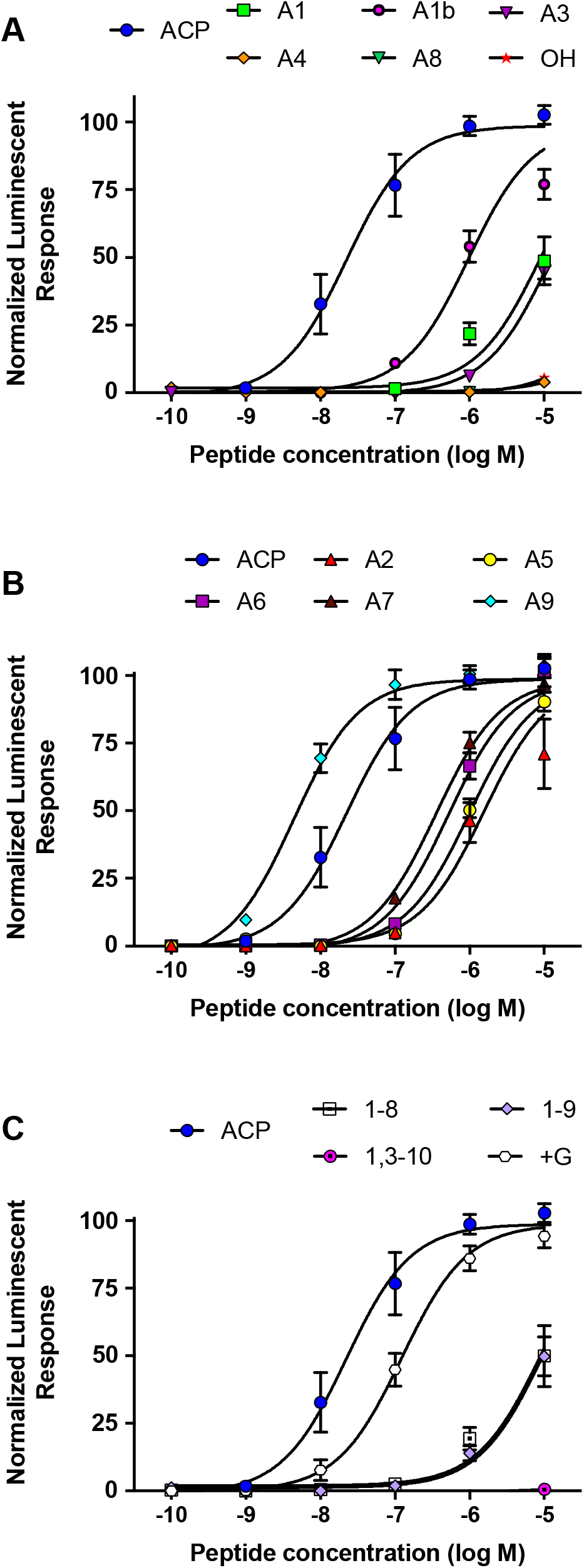
Dose–response curves of the native *A. aegypti* ACP along with synthetic analogs that contain single substitutions or other modifications, which were tested for their activity on the ACP receptor (ACPR) using a cell-based bioluminescent assay. (A) Single amino acid substitutions with alanine or modifications to the normally blocked N- or C-termini resulting in a significant reduction to their activity on the ACPR. (B) Single amino acid substitutions with alanine to the native ACP sequence having minimal effects to the activity of the tested analogs indicating these substitutions are well tolerated. (C) Analogs with internal or terminal truncations or extension (i.e. +Gly analog) relative to the native ACP sequence conferring significant reductions to their activity on the ACPR. The half maximal effective concentration (EC_50_) for each analog along with the corresponding change in activity relative to native *A. aegypti* ACP is provided in Table 1. Luminescence is plotted relative to the maximal response achieved when 10^−5^M Aedae-ACP was applied to the ACPR. Data represent mean +/− standard error of three independent biological replicates.

### Activity of natural arthropod AKH analogs on mosquito AKHR-IA

Taking advantage of the variety of natural AKH analogs in arthropods, we examined a subset of insect AKH peptides (see Table 2) with unique substitutions compared to the endogenous mosquito AKH peptide and determined their activity on the AKH receptor, *A. aegypti* AKHR-IA. From this analysis, the least effective analog tested was the C-terminally truncated peptide, Lacol-AKH-7mer, whose activity was nearly completely abolished with only ~7% AKHR-IA activation at the highest tested concentration (10μM), which represents a reduced efficacy of at least five orders of magnitude (Figure 2A). The next least active AKH analog was the cricket peptide, Grybi-AKH, which has five residues substituted over the length of the octapeptide compared to the mosquito AKH (see Table 2). Specifically, leucine and threonine in positions two and three are replaced by valine and asparagine, respectively. Further, threonine, proline and serine in positions five to seven are replaced with serine, threonine and glycine, respectively. As a result of these combined substitutions, Grybi-AKH activity on the *A. aegypti* AKHR-IA was compromised by over three orders of magnitude (1032-fold reduced activity) compared to the endogenous mosquito AKH (Table 2 and Figure 2A). We next tested the AKH from the dragonfly, *Libellula auripennis*, which allowed us to isolate a subset of the substitutions of Grybi-AKH, specifically positions two and three alone. The activity of Libau-AKH was improved nearly two-fold over Grybi-AKH; however, the activity of this peptide on the *A. aegypti* AKHR-IA was still reduced by 548-fold compared to Aedae-AKH (Table 2). The next AKH analog examined was that from the firebug *Pyrrhocoris apterus* (Pyrap-AKH), which has two substitutions in positions three and seven, both asparagine, for the natural threonine and serine, respectively, found in *A. aegypti* AKH. The activity of Pyrap-AKH was found to be improved compared to Libau-AKH and Grybi-AKH by 1.6 and 3-fold, respectively. However, these substitutions resulted in the activity of Pyrap-AKH on the *A. aegypti* AKHR-IA being compromised by 337-fold relative to the native AKH (Figure 2A). To better understand the importance of N-terminal residues, we next tested a second AKH from Odonata, this analog from *Erythemis simplicicollis* (Erysi-AKH) where the second position valine found in Libau-AKH is instead leucine, which is conserved with the *A. aegytpi* AKH. Thus, the activity of this AKH analog having only a single amino acid substitution in the third position (threonine to asparagine; see Table 2) resulted in Erysi-AKH eliciting activity on the *A. aegypti* AKHR-IA which was 310-fold less compared to native mosquito AKH (Figure 2A). When testing an AKH analog having a single equivalent substitution in only position seven (serine to asparagine), as is naturally available in the American cockroach peptide Peram-CAH-II, this substitution was completely tolerated since effectively no difference in the activity of this analog on the AKHR-IA relative to the *A. aegypti* AKH was detected (Figure 2B). Indeed, we could next verify the criticalness of the third position threonine residue by examining the activity of another cockroach AKH analog, V^2^-Peram-CAH-II (a synthetic analog and not naturally occuring in *P. americana*), which has a second position substitution of valine for leucine along with the seventh position substitution of asparagine for serine (Table 2). The activity of V2-Peram-CAH-II was found to be much improved, albeit still 69-fold less effective on the AKHR-IA compared to the native *A. aegypti* AKH (Figure 2B). Since it was demonstrated that the C-terminal tryptophan was fundamental for bioactivty, with truncated analogs having little or no activity (see above for Lacol-AKH-7mer), we were interested to examine if extension of the C-terminus may also equally impact the activity of the AKH analog on the mosquito *A. aegypti* AKHR-IA. To do this, we utilized the AKH analog from the domestic silkmoth, *Bombyx mori* (Bommo-AKH), which contains an extended C-terminus comprised of a glycine and glutamine residue (resulting in a decapeptide). Additionally, Bommo-AKH has a seventh position substitution of glycine for serine; however, considering the extension on the C-terminus as well as the substitution in position seven, this analog was surprinsingly quite tolerated with only 11-fold reduced activity on the *A. aegypti* AKHR-IA (Figure 2B). Finally, examining naturally occuring analogs of this peptide family containing single substitutions in the C-terminal region, but sparing the eighth critical tryptophan residue, the findings demonstrate that these changes are quite tolerated. It was noted above that the cockroach AKH, Peram-CAH-II, which has a seventh position serine substituted with asparagine, had near identical efficacy to the native mosquito AKH. Similarly, an AKH analog from the sphingid moth, *Hippotion eson* (Hipes-AKH-I), which has a serine substitution for proline in the sixth position, also was well tolerated with only a 8-fold reduced activity on the *A. aegypti* AKHR-IA (Figure 2B). Lastly, using the AKH analog from the black horse fly, *Tabanus atratus* (Tabat-AKH), the results demonstrated that another substitution in the seventh position, in this case a glycine for the naturally occurring serine in Aedae-AKH, is indeed well tolerated since the activity of this analog matched and partially exceeded (~30% improved efficacy) the activity of the native mosquito AKH (see Table 2; Figure 2B).

**Figure 2.**
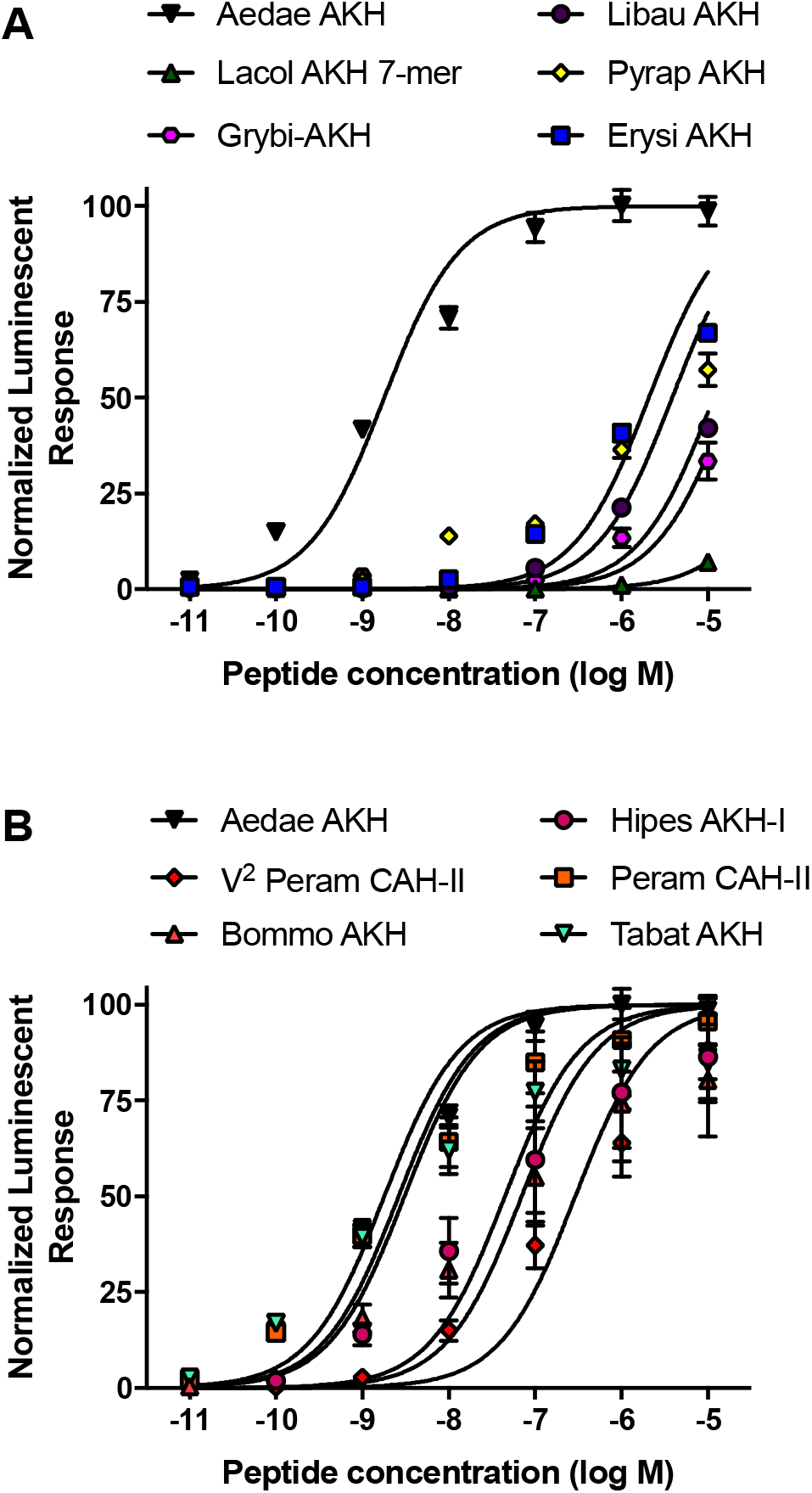
Dose–response curves of the native *A. aegypti* AKH along with mostly naturally occurring analogs from a variety of insects that contain substitutions or other modifications, which were tested for their activity on the AKHR-IA receptor using a cell-based bioluminescent assay. (A) Naturally occurring insect AKH analogs having critical substitutions that are not well tolerated by the AKHR-IA, with moderately compromised activity relative to the native *A. aegypti* AKH. (B) Naturally occurring insect AKH analogs, or a synthetic analog (V^2^-Peram-CAH-II), having substitutions that are well tolerated by the AKHR-IA, with marginally compromised or slightly improved activity relative to the native *A. aegypti* AKH. The half maximal effective concentration (EC_50_) for each analog along with the corresponding change in activity relative to native *A. aegypti* AKH is provided in Table 2. Luminescence is plotted relative to the maximal response achieved when 10^−5^M Aedae-AKH was applied to the AKHR-IA. Data represent mean +/− standard error of three independent biological replicates.

## Discussion

In light of the high fidelity of the three GnRH-related systems (Hansen et al., 2010; Oryan et al., 2018; Wahedi and Paluzzi, 2018), the current study set out to examine for the first time the ligand structure-activity relationship for an insect ACP receptor. This is of utmost importance in order to gain insight on the specific structural features of the ACP system given that extensive studies have been carried out previously on AKH receptors in a variety of species using natural and synthetic analogs of this neuropeptide family (Caers et al., 2012; Fox and Reynolds, 1991; Gäde et al., 2000; Gäde, 1992; Gäde, 1993; Gäde et al., 2016; Keeley et al., 1991; Lee and Goldsworthy, 1996; Lee et al., 1996; Lee et al., 1997; Marco and Gäde, 2015; Marco and Gäde, 2019b; Poulos et al., 1994; Velentza et al., 2000; Ziegler et al., 1991; Ziegler et al., 1998). By examining the activity of a variety of ACP analogs on the activation of the *A. aegytpi* ACP receptor (ACPR), including single alanine substitutions along with truncated or extended analogs, a few observations can be highlighted. Firstly, the overall charge of the peptide does not appear to play a role with regards to its influence on receptor activation since substitution of the basic (arginine) or acidic (aspartic acid) residues did not lead to a highly detrimental effect. Specifically, one would assume that the ACP receptor may prefer a neutral ligand given that the native ACP has no net charge (i.e. is neutral) since it has both a single basic and acidic residue. However, the modified analogs which replace the basic (position 6) or acidic (position 7) residues with alanine, create negatively or positively charged molecules, respectively, with both being quite active and so neither proved detrimental substitutions. Secondly, with the exception of the aromatic tryptophan residue in the eighth position, the C-terminal residues of ACP do not appear to be highly important since alanine substitutions in positions five through seven only marginally impacted activity between ~16 to 45-fold whereas alanine substitution in position nine resulted in this analog having five-fold increased activity. Amidation of the C-terminal residue was critical since the free acid analog exhibited no activity; however, it is noteworthy that the C-terminally extended peptide having an amidated glycine was also quite effective with ACPR activation compromised by only 6-fold. Previous studies examining free acid analogs of AKH (including both *in vitro* and *in vivo* assays) have demonstrated that in some cases the absence of a blocked C-terminus is less critical (Caers et al., 2012; Caers et al., 2016; Lee et al., 1996), whereas other investigations have revealed the presence of an amidated C-terminus to be very important with poor or no responses with analogs lacking this feature (Gäde and Hayes, 1995; Gäde, 1990). These observed differences in response to key structural elements, including the normally amidated C-terminus of AKH, may reflect the occurrence of inter- and intra-species receptor subtypes which may be differentially sensitive to these modifications (Caers et al., 2016; Gäde and Hayes, 1995), and could also point to the well documented phenomenon that some species have AKH receptors that are more promiscuous whereas others are very strict in the structural characteristics of their particular ligand (Caers et al., 2012; Gäde, 1990; Marchal et al., 2018). This latter scenario is more in-line with observations for the *A. aegypti* ACP system where only a single functional receptor variant occurs (Wahedi and Paluzzi, 2018), which we show herein does not tolerate the absence of the C-terminal amidation and is thus of high importance for the ACP system. Thirdly, both aromatic residues were essential for activity on the ACPR while N-terminal residues of the peptide, including mainly threonine and pyroglutamic acid, were also highly critical since alanine substitutions were not tolerated leading to significantly reduced activity of these analogs relative to the native ACP peptide. However, alternatively blocking this alanine substitution on the N-terminus through acetylation improved the activity of this analog by 10-fold. Similar observations have been made with AKH analogs tested using both *in vivo* and *in vitro* assays whereby complete removal of the N-terminal pyroglutamate abolishes activity of the AKH analog whereas the alternatively blocked N-terminus (i.e. N-Acetyl-Ala), or even simply glutamate or glutamine that can spontantenously undergo cyclization (Schilling et al., 2008), were shown to have improved activity, albeit lower than the native AKH (Caers et al., 2016; Gäde and Hayes, 1995; Marco and Gäde, 2019b). Lastly, although alanine substitution for valine in the second position led to a relatively minor effect on ACPR activation (~70-fold reduced activity), the internally truncated analog omitting valine within the N-terminal region elicited no activation of the ACPR, indicating the spacing between the more critical residues, namely pyroglutamate and polar threonine, is essential for ACP peptide activity on the ACPR. Thus, perhaps in common with AKH structural features, the switch between adjacent hydrophobic and hydrophilic residues is necessary in order to properly ‘fit’ and bind with its particular receptor (Gäde and Hayes, 1995) so that not only is the ACP analog shorter by one amino acid (a nonapeptide), but moreover, the entire structural configuration of the peptide is disrupted and the whole sequence is no longer in sync with its receptor. This corroborates with studies that have examined the conservation of the ACP primary structures across insects from different orders (including Diptera, Lepidoptera, Coleoptera, Hymenoptera and Hemiptera) as well as from various species within the same order (seventeen species of Coleoptera), where the N-terminal pentamer sequence, when the ACP system is present, is nearly completely conserved across these various species with a consensus motif of pQVTFS-, whereas significant sequence variability occurs within the C-terminus of the ACP sequence with deca-, nona- and even dodecapeptides reported (Hansen et al., 2010; Veenstra, 2019).

Considering the resultant activation of the mosquito AKH receptor in response to the various mostly naturally-occurring insect AKH analogs examined in the current study, these findings indicate that these bioanalogs can be grouped into distinct categories based on their determined potencies on the *A. aegypti* AKHR-IA. The first group includes natural or synthetic AKH analogs having substitutions in critical positions, including the absence of the C-terminal tryptophan that occurs in the heptapeptide, Lacol-AKH-7mer, which is not tolerated whatsoever since activity is completely abolished and is therefore of greatest importance for AKH receptor activation. Observations correlating with these findings have been made in various studies utilizing *in vitro* heterologous assays as well as *in vivo* bioassays monitoring lipid- and/or carbohydrate-mobilizing actions of AKH analogs. Specifically, removal or replacement of the critical aromatics that are archetypal features of AKH neuropeptides found in positions four and eight demonstrates the absolute requirement of these features for receptor activation in heterologous systems or *in vivo* biological activity (Gäde, 1992; Gäde, 1993; Marco and Gäde, 2015; Marco and Gäde, 2019b).

The second group in the current study includes analogs that were similar, but had a higher efficacy, to the tested C-terminally truncated peptide lacking the eighth position tryptophan. In particular, additional residues of major importance were identified in both the amino and carboxyl region of the octapeptide, since Grybi-AKH, which has substitions in positions 2-3 as well as 5-7 compared to the native *A. aegypti* AKH, demonstrated a large reduction in activity by over three orders of magnitude. Interestingly, this implies that despite Grybi-AKH retaining the prototypical features of an AKH, including the pyroglutamic acid in the first position, phenylalanine in position four, tryptophan in position eight and an amidated C-terminus, the *A. aegypti* AKH receptor is quite specific compared to other insect AKH receptors including, for example, the locust, *Schistocerca gregaria* or the pea aphid, *Acyrthosiphon pisum*, whose receptors responded similarly to both the mosquito AKH and Grybi-AKH (Marchal et al., 2018). Similar findings were reported for the AKH receptor from another dipteran, namely the fruit fly *D. melanogaster*, which is also highly selective and was not strongly responsive to non-dipteran AKH analogs (Marchal et al., 2018). In order to determine whether the substitutions in positions 2-3 versus 5-7 were of greatest importance, we examined the activity of a natural AKH analog from *L. auripennis* (Libau-AKH), whose sequence matches that of native *A. aegypti* AKH except for positions 2-3 (valine and asparagine) that are instead shared with Grybi-AKH. This allowed us to identify that the substitutions in this N-terminal region are much more critical and are not well tolerated by the AKH receptor system in *A. aegypti* since this analog diplayed only a marginal improvement (i.e. nearly two-fold) compared to Grybi-AKH. This finding is corroborated by recent analysis of select dipteran AKH analogs on the *A. aegypti* AKH receptor, which showed that amino acid changes at positions five (serine in both *Glossina morsitans* AKHs and in *D. melanogaster* AKH compared to threonine in *A. aegypti* AKH) or seven (glycine in *G. morsitans* AKH-I and aspartic acid in *D. melanogaster* AKH compared to serine in *A. aegypti* AKH) are indeed well tolerated with these dipteran AKH analogs being similarly active to the endogenous mosquito AKH (Marchal et al., 2018). It was also previously shown in other dipteran AKH systems where structure-activity relationships were investigated, that the least tolerated substitutions were localized to the N-terminal region, with substitutions to residues in positions two through five from the N-terminus proving crucial for activation of *D. melanogaster* and *A. gambiae* AKH receptors, which the authors described was due to this core region of the peptide forming a predicted β-strand necessary for receptor interaction (Caers et al., 2012). Comparatively, although only fruit fly AKH analogs were tested, substitutions to position seven alone and positions six plus seven for the *D. melanogaster* and *A. gambiae* AKH receptors, respectively, were far more tolerated with little or no influence on the activation of the corresponding receptor (Caers et al., 2012). This supports our conclusions that the C-terminal region of the AKH octapeptide, excluding the critical tryptophan in the eighth position, is not as critical for activity. As a putative reason for this, it is argued that the side chains of a subset of these amino acids may be buried in the assumed β-turn formed by the residues in the C-terminal region of AKH neuropeptides (Caers et al., 2012; Gäde, 1992; Gäde and Hayes, 1995), which is now supported by molecular modeling and nuclear magnetic resonance spectroscopy studies that have confirmed the β-turn configuration in the AKH secondary structure (Jackson et al., 2014; Nair et al., 2000; Nair et al., 2001; Zubrzycki and Gäde, 1994).

Interestingly, structure-activity studies on the two tsetse fly (*G. morsitans*) AKH receptors using synthetic analogs of the pair of endogenous AKHs identified the most critical sites being the aromatic residues in positions four and eight, but the next most important features included positions 1-3 from the N-terminus with activity of these analogs eliciting only 20-50% activity compared to the two endogenous AKH peptides (Caers et al., 2016). On the other hand, the region of the *G. morsitans* AKH diplaying increased permissiveness for substitutions included positions six and seven, where glycine substitutions of these sites resulted in these analogs having between 65-97% activity compared to the efficacy of the native AKHs. Interestingly, however, substitutions in the fifth position of the octapeptide were not tolerated, since activity of these analogs was reduced to only 11-13% that of the natural AKHs, which nearly matches the effect of substitutions in the aromatic residue sites at positions four and eight (Caers et al., 2016). Supporting the notion that substitutions in positions 5-7 are more permissive in the mosquito AKH receptor system, we examined the activity of two additional natural AKH analogs, which in comparison to *A. aegypti* AKH, have substitutions in either the third position alone or combined with a substitution of the seventh position residue. In particular, the firebug *P. apterus* AKH (Pyrap-AKH) has the third position threonine found in *A. aegypti* AKH replaced with asparagine while the seventh position serine is also replaced by asparagine, resulting with this analog having ~337-fold reduced activity relative to the native mosquito AKH. Next, by comparing the tolerance of these two substitutions in Pyrap-AKH with that occurring in the AKH from *E. simplicicollis* (Erysi-AKH), which has only a single substitution in the third position with threonine replaced by asparagine, reveals this AKH analog has similar activity as Pyrap-AKH with ~310-fold reduced activity relative to the endogenous *A. aegypti* AKH.

The third group includes analogs with activities similar to, or in some cases, stronger than the endogenous *A. aegypti* AKH that contained substitutions or insertions in the C-terminal region, while sparing the critical tryptophan residue. Specifically, we next tested two natural AKH analogs with substitutions in the C-terminal region of the peptide, maintaining the core 5-6 residues on the N-terminus. Specifically, the domestic silkmoth AKH (Bommo-AKH), has a glycine substitution in an equivalent location to the serine in position seven and also has an extended C-terminus containing glycine and glutamine that follows tryptophan in the eighth position, forming a decapeptide. Despite these changes within the C-terminal region, this AKH analog retained strong activity on the *A. aegypti* AKHR-IA. Similarly, analysis of Hipes-AKH-I from the hawk moth, *Hippotion eson*, which along with another member of the genus, each produce the greatest number of AKH analogs so far reported from a single insect (Gäde et al., 2013), revealed this analog to be very active, with only a marginal reduction in activity on the mosquito AKHR-IA. Notably, Hipes-AKH-I shares an identical sequence to the Lacol-AKH-7mer with the one exception that the latter lacks the terminal tryptophan residue. This indicates that serine for proline substitution in the sixth position is in fact well tolerated and is not the root cause of the abolished activity of the Lacol-AKH-7mer, which instead is most certainly due to the absence of the C-terminal amphipathic tryptophan residue with its large indole side chain.

Considering the resulting activity of these various AKH analogs, the findings support that substitutions in positions 5-7 are more permissive, whereas comparatively, substitutions in positions 2-3 are not well tolerated by the *A. aegypti* AKH receptor system. Therefore, given the importance of substitutions in positions 2-3, we aimed to discern which of these two sites were the most critical for eliciting AKHR-IA functional activation. To achieve this aim, we tested a synthetic analog of a cockroach *P. americana* AKH, V^2^-PeramCAH-II, which has a second position valine substituted for the naturally occurring leucine in *A. aegypti* AKH but also contains a substitution in the seventh position with asparagine replacing serine. The results from analysis of this analog confirm the valine substitution for leucine in the second position is far better tolerated compared to the asparagine substitution for threonine in the third position, since V^2^-Peram-CAH-II displayed nearly five-fold improved activity on the *A. aegypti* AKHR-IA compared to Pyrap-AKH (which has asparagine substitutions in both positions three and seven). Thus, although both of these residues are important for functional activation of the mosquito AKHR-IA, substitutions of the second position proved this site to be the most critical. One additional matter to consider is the seventh position substitution, which is an asparagine in Pyrap-AKH, V^2^-Peram-CAH-II and is the only substitution in the natural AKH analog, Peram-CAH-II. The activity of Peram-CAH-II was equivalent to the native mosquito AKH, which confirms that this seventh position is quite permissive to substitutions and, more importantly, that the markedly reduced activity of the AKH analogs Pyrap-AKH and V^2^-Peram-CAH-II is solely a result of the substitutions in the second and third positions. Finally, to our surprise, the survey of natural AKH analogs revealed that Tabat-AKH, which has a single glycine substitution in position seven replacing the natural serine of *A. aegypti* AKH, may serve as a potential lead substance for design of a superagonist since this natural AKH analog elicited 30% improved activity relative to the native mosquito AKH. It appears that the more flexible and non-polar Gly residue allows the ligand a tighter binding to the receptor than the polar and neutral Ser. This is particularly interesting given the permissiveness of substitutions in the vicinity of the C-terminus (excluding the eighth position tryptophan) established in the current study and previously for the *A. aegypti* AKHR-IA (Marchal et al., 2018) as well as homologous receptors from other dipterans (Caers et al., 2012; Marchal et al., 2018), which may allow a greater variety of lead compounds to be designed and tested that could interfere with this neuropeptide system and perturb its normal functioning vital for mobilizing energy substrates in insects.

Integrating the novel data from the present study analyzing structural characteristics of the two most recently diverged GnRH-related family members in arthropods, we can infer with some degree of confidence how these ligands act only upon their respective receptors. The ACP sequence in many insect species often contain charged residues in positions 6, 7 or 9, which in instances where only a single basic or acidic residue occurs, provide an overall charge to the peptide (Hansen et al., 2010; Veenstra, 2019). However, our results revealed that the overall charge of the ACP analog did not largely compromise activity since substitutions of these charged residues with alanine were quite tolerated by the ACPR. Comparatively, AKH receptors do not like charges associated with their ligands since no basic residues occur and only a few natural AKH family members containg an acidic residue (e.g. aspartic acid at position 7), which have been shown to be not well tolerated by AKH receptors in *L. migratoria* and *P. americana* (Gäde, 1991; Gäde, 1993; Gäde et al., 1990). Another feature differentiating the AKH and ACP peptides in *A. aegypti* is the presence of an alternating pattern of hydrophobic and hydrophilic residues over the entire length of the AKH, which is shared with other AKH family members (Gäde and Hayes, 1995), whereas the ACP contains a motif of three hydrophilic residues at its core residues 5-7, which may lead to incompatability with the AKHR-IA. Additionally, clearly the valine substitution in position 2 is not tolerated well by the *A. aegypti* AKHR-IA, and so this residue which is present in position 2 of the ACP sequence may also contribute towards the inactivity of this peptide on the AKHR-IA. On the other hand, since the AKH peptide from *B. mori* (a decapeptide) was quite active on the *A. aegypti* AKHR-IA, the longer length of ACP is unlikely to be the cause of its inactivity on the AKHR-IA.

Our findings also set the framework towards understanding why the *A. aegypti* AKH has no activity on the ACP receptor. As we saw with the AKHR-IA, substitution of valine for leucine in position 2 led to nearly a 70-fold reduction in receptor activity. On the other hand, the removal of valine and its replacement with alanine in the ACP sequence led to a similar 70-fold reduction in activity on the ACPR. Thus, the presence of valine in position two is needed for optimal activity for ACPR whereas its absence in the same position (but on the AKH peptide) is necessary for optimal activation of AKHR-IA. Again, as discussed earlier, the importance of charged residues for optimal ACPR activity that are often found in insect ACPs (Hansen et al., 2010; Veenstra, 2019), whereas charged residues are unusually found on AKH peptides and are not well tolerated by AKH receptors that have been studied either *in vivo* or *in vitro* (Gäde, 1991; Gäde, 1993; Gäde et al., 1990). Finally, one clear finding from these studies is that ACP analogs that are too short (nonapeptide or octapeptide), even if they contain all the hallmark features, as demonstrated by the C-terminally truncated analogs for example, fail to activate ACPR. Thus, the length of the ligand is most relevant and the binding pocket of ACPR requires at least a decapeptide leaving the *A. aegypti* AKH, an octapeptide, too short for ACPR binding and activation.

## Author Contributions

GG and JP designed the synthetic analogs and wrote the manuscript. AW performed all the experiments and AW, JP and GG analyzed the data. All authors have contributed towards the revisions and have granted approval of the final manuscript submitted for publication.

## Funding

This work was funded by a Natural Sciences and Engineering Research Council of Canada (NSERC) Discovery Grant and an Ontario Ministry of Research, Innovation and Science Early Researcher Award to JP and Incentive Grant from the National Research Foundation (Pretoria, South Africa; grant number 85768 [IFR13020116790] to GG) and by the University of Cape Town (block grant to GG).

## Conflict of Interest Statement

The authors declare that the research was conducted in the absence of any commercial or financial relationships that could be construed as a potential conflict of interest.

